## Supplementay Figures for "Promoter-proximal elongation regulates transcription in archaea"

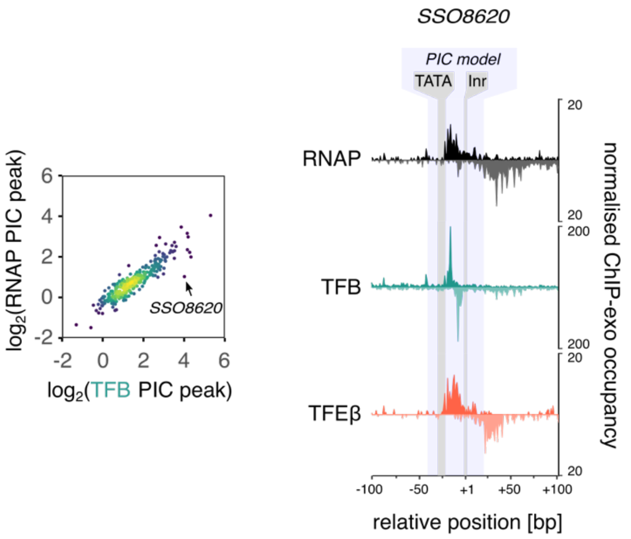


**Supplementary Figure 1: Lower RNAP recruitment is associated with narrower TFB ChIP-exo footprints**

ChIP-exo profile of the *SSO8620* promoter that shows low RNAP and TFEβ ChIP-exo signal relative to TFB. The left panels show the quantifications for RNAP and TFB PIC signal as in Figure 1d with the position of *SSO8620* indicated. The right panel shows the ChIP-exo signal for RNAP, TFB, and TFEβ. RNAP and TFEβ profiles show similar broad footprints as for the aggregate profiles in Figure 1b that can be attributed to PICs. In contrast, the TFB profile is dominated by a sharper footprint that can be attributed to ternary TFB-TBP-DNA complexes lacking RNAP and TFEβ. All ChIP-exo data represent the geometrical mean of three biological replicates.

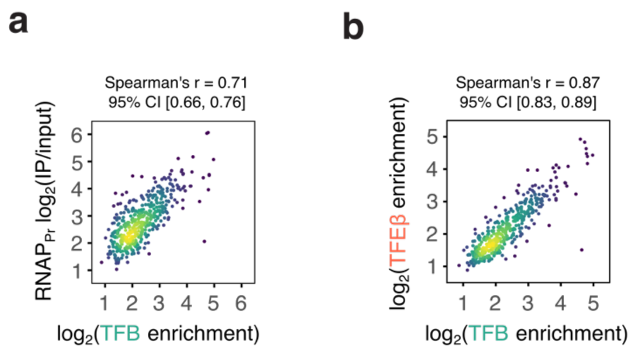


**Supplementary Figure 2: ChIP-seq data are consistent with uniform PIC assembly during exponential growth**

(a) and (b) Scatter plots depicting correlation of TFB promoter occupancy with RNAP (a) and TFEβ promoter occupancy (b) under exponential growth conditions based on ChIP-seq data (n=454 promoters). Data show the geometric mean of two biological replicates. These data are overall consistent with ChIP-exo data (Figure 1de). Because of the limited resolution of ChIP-seq occupancy data, the RNAP promoter occupancy quantification ($RNAP_{Pr}$) comprises RNAP present in PICs as well as promoter-proximal TECs. Therefore the correlation of $RNAP_{Pr}$ with with TFB is lower than the corresponding ChIP-exo signal correlation (Figure 1d) or the correlation between TFB and TFEβ which are both specific to PICs (panel b).

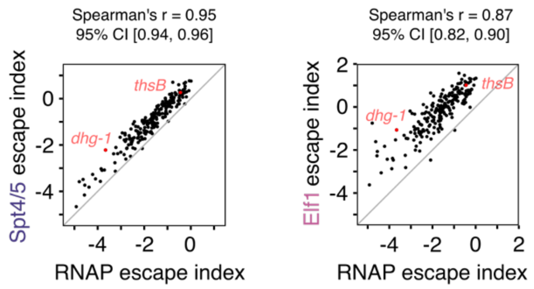


**Supplementary Figure 3: Correlated promoter-proximal accumulation of elongation factors and RNAP**

Scatter plots depicting the correlation between escape indices calculated for RNAP and elongation factors Spt4/5 (left panel) and Elf1 (right panel) for 212 TUs. Escape indices represent the mean of two biological replicates.

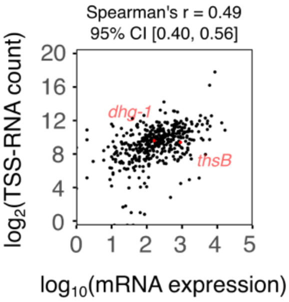


**Supplementary Figure 4: Correlation between TSS-RNAs and mRNA levels**

Scatter plot depicting the correlation between mRNA expression levels and TSS-RNA (n=438 TUs). TSS-RNA data represent the geometric mean of two biological replicates. Rockhopper [75] estimates of mRNA levels are based on two biological replicates.

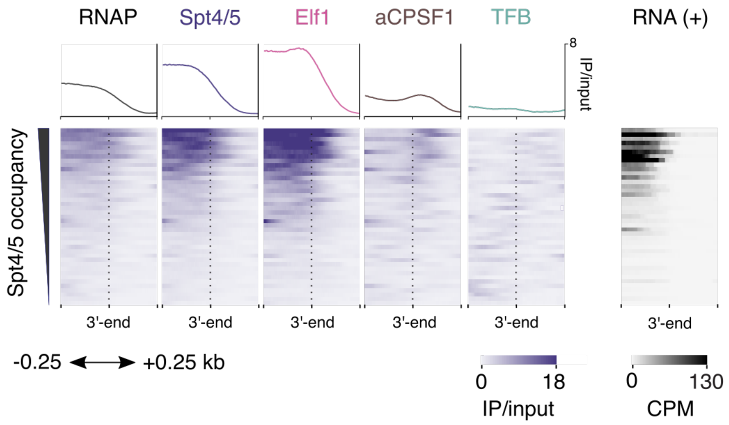


**Supplementary Figure 5: aCPSF1 association with transcription termination sites.** Heatmap of RNAP, elongation factors Spt4/5 and Elf1 and termination factor aCPSF1 ChIP-seq occupancy at 41 predicted RNA 3′-ends during exponential growth. Data represent one representative of two biological replicates. RNA 3′-ends as putative termination sites were predicted based on RNA-seq data and filtered for the expected Spt4/5 occupancy decay downstream of termination sites as well as the absence of TFB peaks to ensure the absence of active promoters and potentially promoter-proximal aCPSF1 peaks in the plotted region. TFB occupancy is depicted as control. The corresponding RNA-seq data for the plus strand are depicted on the right.

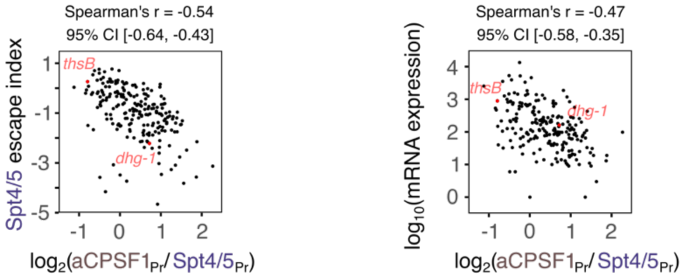


**Supplementary Figure 6: aCPSF1 load on Spt4/5-containing TECs is negatively correlated to TEC escape**

The left panel shows a scatter plots depicting the negative correlation between Spt4/5 escape index (mean of two biological replicates) and the relative load of aCPSF1 on the Spt4/5-bound TEC calculated as $aCPSF1_{Pr}$ to $Spt4/5_{Pr}$ ratio (geometric mean of two biological replicates, n=212 TUs). The right panel shows the correlation of aCPSF1 load calculated relative to $Spt4/5_{Pr}$ replicates (geometric mean of two biological replicates, n=211 TUs) with mRNA expression levels. Rockhopper [75] estimates of mRNA levels are based on two biological replicates.

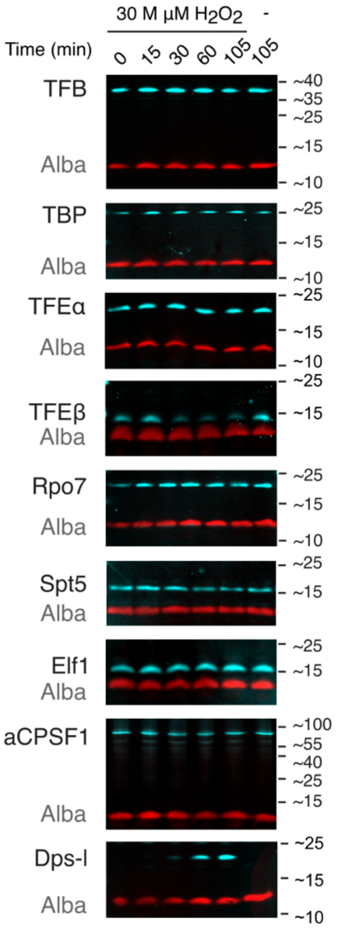


**Supplementary Figure 7: TFEβ depletion in response to oxidative stress**

Multiplex immunodetection of initiation factors (TFB, TBP, TFEα, TFEβ) RNAP subunits Rpo4/7, elongation factors (Spt4/5, ELf1) and termination factor aCPSF1 during a time course after $H_2O_2$-treatment (cyan signal). Detection of the oxidative stress marker protein Dps-l is shown as control. The stably expressed chromatin Alba (red signal) was used as loading control. A representative blot from three biological replicates is shown.

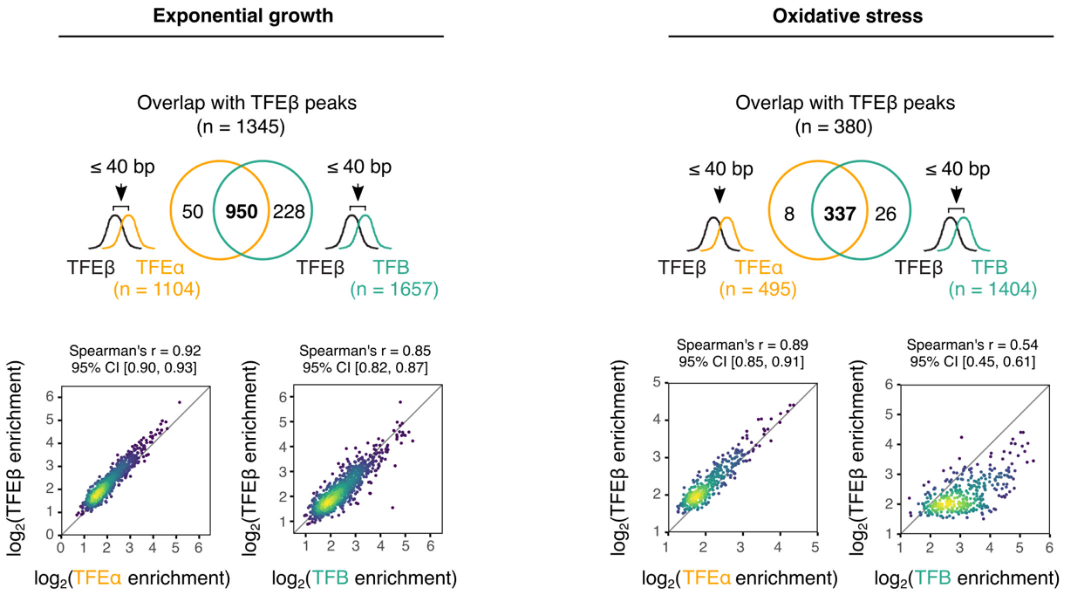


**Supplementary Figure 8: Attenuated recruitment of TFE to PICs under oxidative stress conditions**

Scatter plots showing the genome-wide correlated enrichment for TFEβ peaks overlapping with both TFEα and TFB under exponential growth (left panel) and oxidative stress conditions (right panel). Under oxidative stress conditions, TFEβ remain strongly correlated to TFEα and both factors show weaker enrichment compared to TFB. Data represent the geometric mean of two biological replicates.

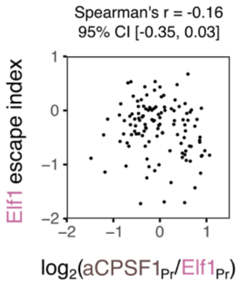


**Supplementary Figure 9: aCPSF1 load on the TEC shows a weaker link to TEC escape under oxidative stress conditions**

Scatter plot depicting the relation between the Elf1 escape index (mean of two biological replicates) and aCPSF1-load calculated as aCPSF1$_{Pr}$ to Elf1$_{Pr}$ ratio (geometric mean of two biological replicates, n=118 TUs). No significant correlation was detected.

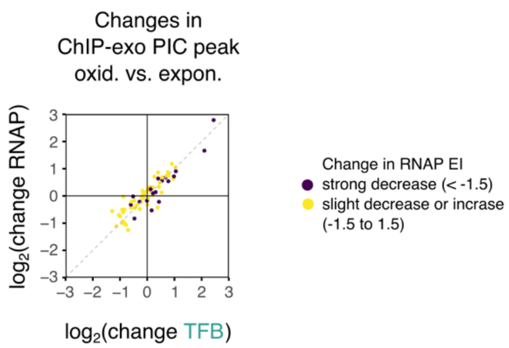


**Supplementary Figure 10: TFB and RNAP PIC occupancy shows equal changes between growth conditions.**

Scatter plot showing $log_2$-fold changes in ChIP-exo PIC signal between oxidative stress and exponential growth phase. PIC signal was calculated as ChIP-exo signal on the non-template strand over a window from -30 to +20 relative to TSS (Figure 1c). Promoters with both mapped TSS [73] and TSS prediction only based on start codon position and ChIP-exo profiles (see methods) were included (n=70). We centred the data by subtracting the mean value for all selected promoters. The promoters of TUs undergoing a larger reduction in RNAP escape index (< 1.5) under oxidative stress conditions compared to exponential growth are labelled in dark purple (n=19). All ChIP-exo quantifications were based on the geometric mean of three biological replicates.

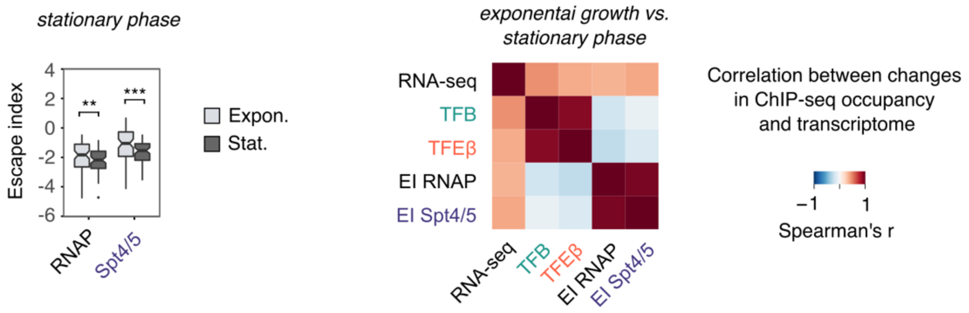


**Supplementary Figure 11: TEC escape and transcriptome changes between exponential growth and stationary phase**

The left panel shows reduced escape of RNAP and Spt4/5 during stationary phase. Boxplots comparing escape indices under exponential growth and stationary phase conditions for TUs accessible for analysis in both conditions (n=74). Differences in escape index distribution were assessed using one-sided paired wilcoxon rank-sum test, *** denotes $p < 0.001$, ** $p < 0.01$.

The right panel depicts a heatmap showing correlated changes in initiation factor occupancy, escape indices and RNA output between exponential growth and stationary phase. Spearman rank correlations were calculated for protein-encoding TUs accessible for analysis in both conditions (n=73). Correlations were calculated from mean escape index and the geometric mean of all other values for two biological replicates.

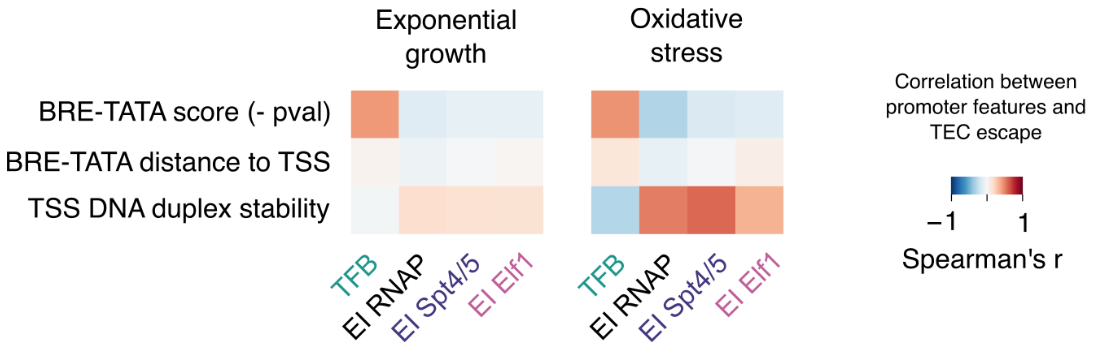


**Supplementary Figure 12: Link between BRE-TATA core promoter elements with TEC escape**

Heatmap depicting the correlation between the archaeal core promoter elements BRE-TATA and TEC escape calculated as escape indices for RNAP, Spt4/5 and Elf1 (mean of two biological replicates). 140 and 93 TUs were include for exponential growth and oxidative stress conditions, respectively. BRE-TATA elements were predicted by MEME for both sets of genes independently and the Spearman correlation was calculated between the motif strength for each promoter and the escape indices. Two controls are included: the positive correlation between TFB promoter occupancy and BRE-TATA score and the positive correlation of TSS DNA duplex stability with RNAP and Spt4/5 escape indices (see Figure 6).

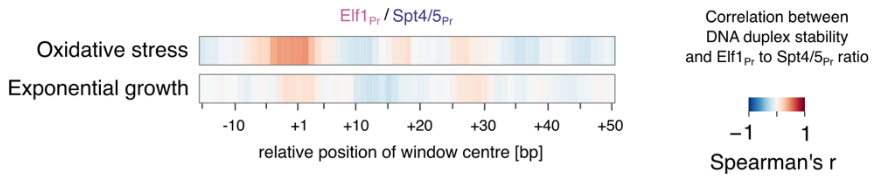


**Supplementary Figure 13: Promoter-proximal TEC assembly is sensitive to DNA duplex stability around the TSS under oxidative stress.**

Heatmaps depicting the correlation between DNA duplex stability and with the ratio of Elf1 to Spt4/5 occupancy in the promoter region (mean of two biological replicates) under oxidative stress (top) and exponential growth conditions (bottom). DNA duplex stability was calculated over a 7 bp sliding window for individual promoters (see methods). Selected TUs with mapped TSS were included (n=93 TUs for oxidative stress and n=140 TUs for exponential growth).

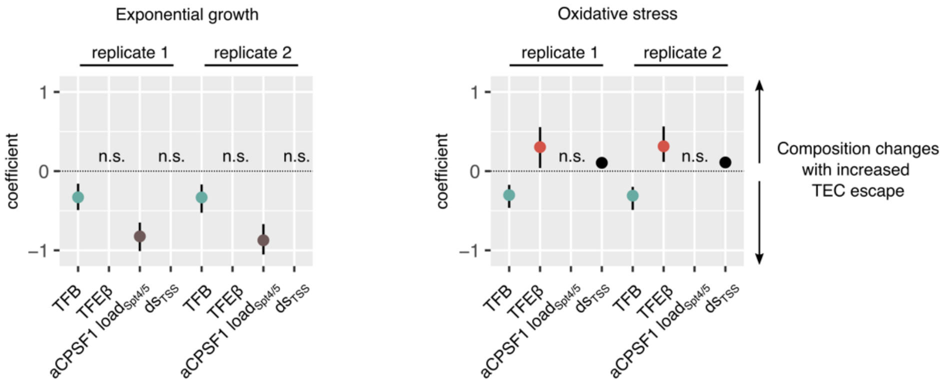


**Likelihood-ratio $\chi^2$-test**

| Explanatory variable | Exponential growth replicate 1 | Exponential growth replicate 2 | Oxidative stress replicate 1 | Oxidative stress replicate 2 |
|---|---|---|---|---|
| TFB | $< 10^{-3}$ | $< 10^{-3}$ | $< 10^{-3}$ | $< 10^{-4}$ |
| TFEβ | n.s. | n.s. | $< 10^{-2}$ | $< 10^{-2}$ |
| aCPSF1 $load_{Spt4/5}$ | $< 10^{-14}$ | $< 10^{-14}$ | n.s. | n.s. |
| $ds_{TSS}$ | n.s. | n.s. | $< 10^{-4}$ | $< 10^{-5}$ |

**Supplementary Figure 14: Multiple regression analysis of TEC escape under exponential growth and oxidative stress conditions**

We generated negative binomial generalised linear models for TEC escape under exponential growth (left panel, n = 137 TUs) and oxidative stress conditions (right panel, n = 92 TUs) based on Spt4/5 escape indices (see methods). A set of four candidates for explanatory variables was tested: TFB and TFEβ promoter occupancy (log-transformed), aCPSF1 load on promoter-proximal TECs (log-transformed aCPSF1$_{Pr}$ to Spt4/5$_{Pr}$ ratio), and TSS DNA duplex stability. Explanatory variables were included in the final models if they improved the models significantly as determined by the likelihood ratio chi-squared test as depicted in the lower panel (threshold for inclusion of $p < 0.05$). "n.s." marks explanatory variables not passing the likelihood ratio test for the model. The models were consistent between biological replicates. Coefficient estimates for two biological replicates are shown. The models were validated by bootstrapping the data set and error bars represent bootstrapped 95% confidence intervals.
